# Acetobacteraceae in the honey bee gut comprise two distant clades with diverging metabolism and ecological niches

**DOI:** 10.1101/861260

**Authors:** G Bonilla-Rosso, C Paredes Juan, S Das, KM Ellegaard, O Emery, M Garcia-Garcera, N Glover, N Hadadi, JR van der Meer, SAGE class 2017-18, F Tagini, P Engel

## Abstract

Various bacteria of the family Acetobacteraceae are associated with the gut environment of insects. Honey bees harbor two distinct Acetobacteraceae in their gut, Alpha2.1 and Alpha2.2. While Alpha2.1 seems to be a gut specialist, Alpha2.2 is also found in the diet (e.g. royal jelly), the hypopharyngeal glands, and the larvae of honey bees. Here, we combined amplicon and genome sequencing to better understand functional differences associated with the ecology of Alpha2.1 and Alpha2.2. We find that the two phylotypes are differentially distributed along the worker and queen bee gut. Phylogenetic analysis shows that Alpha2.2 is nested within the acetic acid bacteria and consists of two separate sub-lineages, whereas Alpha2.1 belongs to a basal lineage with an unusual GC content for Acetobacteraceae. Gene content analysis revealed major differences in the central carbon and respiratory metabolism between the two phylotypes. While Alpha2.2 encodes two periplasmic dehydrogenases to carry out oxidative fermentation, Alpha2.1 lacks this capability, but instead harbors a diverse set of cytoplasmic dehydrogenases. These differences are accompanied by the loss of the TCA cycle in Alpha2.2, but not in Alpha2.1. We speculate that Alpha2.2 has specialized for fast-resource utilization through incomplete carbohydrate oxidation, giving it an advantage in sugar-rich environments such as royal jelly. On the contrary, the broader metabolic range of Alpha2.1 may provide an advantage in the worker bee hindgut, where competition with other bacteria and flexibility in resource utilization may be relevant for persistence. Our results show that bacteria belonging to the same family may utilize vastly different strategies to colonize niches associated with the animal gut.

## Introduction

According to a recent review on the taxonomy of Alphaproteobacteria (Munoz-Gomez, et al. 2019) and the standardized genome phylogeny-based taxonomy of Parks et al. (Parks, et al. 2018), the family Acetobacteraceae (Rhodospirillales) is comprised of an externally branching acidophilic/neutrophilic group and an internal acetous group. The latter group includes acetic acid bacteria (AABs), which constitute the vast majority of the described taxa of the Acetobacteraceae (Komagata, et al. 2014).

AAB inhabit sugar-rich environments and use a rather exceptional strategy to gain energy. They oxidize sugars or sugar alcohols on the periplasmic side of the cell envelop with the help of membrane-bound dehydrogenases that are linked to the respiratory chain in a process known as oxidative fermentation (Matsushita and Matsutani 2016). This particular oxidative metabolism results in the accumulation of fermentation products (such as acetic acid) in the environment. AABs naturally occur in association with plants, flowers, and fruits (Bartowsky and Henschke 2008; Pedraza 2016; Yamada and Yukphan 2008). They also play key roles in food and beverage fermentations (De Roos and De Vuyst 2018). In addition, AABs are being increasingly described to be associated with different insect species that rely on sugar-based diets, such as fruit flies, mosquitoes, sugarcane leafhoppers, mealybugs, honey bees, and bumble bees (Chouaia, et al. 2014; Crotti, et al. 2010).

Honey bees feed on a highly sugar-rich diet composed of nectar, honey, and pollen (Brodschneider and Crailsheim 2010). While their gut microbiota is relatively simple,most of the commonly found phylotypes are specialized to live in the bee gut environment (Bonilla-Rosso and Engel 2018; Kwong and Moran 2016). These phylotypes are consistently present in adult worker bees worldwide and belong to deep-branching phylogenetic lineages that have so far only been detected in different species of honey bees, bumble bees, and stingless bees (Kwong, et al. 2017). Two of these phylotypes, originally referred to as Alpha2.1 and Alpha2.2, belong to the Acetobacteraceae (Cox-Foster, et al. 2007; Martinson, et al. 2011). Both are frequently detected in adult worker bees. However, in contrast to the so-called core phylotypes of the honey bee gut microbiota *(Gilliamella, Snodgrassella, Lactobacillus* Firm4, *Lactobacillus* Firm5, and *Bifidobacterium),* Alpha2.1 and Alpha2.2 are not present in every individual worker bee and often their relative abundance is low (Kapheim, et al. 2015; Kešnerová, et al. 2019; Kwong, et al. 2017; Martinson, et al. 2011; Martinson, et al. 2012; Moran, et al. 2012; Powell, et al. 2018). Interestingly, they both belong to the few phylotypes that have repeatedly been found in the gut of the honey bee queen, which is the only reproductive female in a honey bee colony (Anderson, et al. 2018; Kapheim, et al. 2015; Powell, et al. 2018).

A recent study showed that Alpha2.2 was predominantly found in the mouth, midgut, and ileum of queens, while Alpha2.1 was more abundant in the rectum (Anderson, et al. 2018). Moreover, Alpha2.2 has also been detected in royal jelly, floral nectar, bee bread (i.e. pollen stores of honey bees), and in bee larvae, indicating that this phylotype can live in more diverse environments than Alpha2.1 (Anderson, et al. 2014; Corby-Harris, et al. 2014; Maes, et al. 2016; Vojvodic, et al. 2013).

Several strains of Alpha2.2 have been isolated, one of which (strain MRM) was described as a novel species, *Bombella apis* (Yun, et al. 2017). Other isolates of Alpha2.2 have been referred to as *Candidatus* Parasaccharibacter apium (Corby-Harris, et al. 2016; Corby-Harris, et al. 2014), but were never formally described as a novel species. Some strains of Alpha2.2 have been suggested to increase larval survival under experimental conditions indicating a possible beneficial role of this phylotype (Corby-Harris, et al. 2014).

Genome analysis of a closely related species, *Bombella intestini*, isolated from the gut of a bumble bee, revealed typical features of other Acetobacteraceae, including the presence of genes for carrying out oxidative fermentation (Li, et al. 2016; Li, et al. 2015). Draft genomes of several strains of Alpha2.2 have been deposited in public databases (Corby-Harris and Anderson 2018), and a comparison with a closely related flower-associated strain, *Saccharibacter floricola,* revealed a number of potentially adaptive changes (Smith and Newton 2018). However, an in-depth functional analysis of the gene content and overall metabolic capabilities of Alpha2.2 relative to Alpha2.1 has not been carried out to date.

Compared to Alpha2.2, much less is known about the phylotype Alpha2.1. Phylogenetic trees based on 16S rRNA gene sequences suggest that Alpha2.1 belongs to a deep-branching lineage within the Acetobacteraceae (Martinson, et al. 2011). The two most closely related species belong to the candidate genus *Commensalibacter* (*Commensalibacter* sp. MX-Monarch01 and *Commensalibacter intestini* A911), isolated from the gut of a butterfly and a fruit fly, respectively (Roh, et al. 2008; Servin-Garciduenas, et al. 2014). Therefore, the phylotype Alpha2.1 is frequently referred to as *Commensalibacter* sp. While genomes have been published for strains of all three *Commensalibacter* species (Kim, et al. 2012; Servin-Garciduenas, et al. 2014; Siozios, et al. 2019), little is known about their gene content, metabolic capabilities, and phylogenetic positioning in respect to other Acetobacteraceae.

The presence of two phylogenetically related phylotypes from a family known to be optimized for fast growth in carbohydrate-rich environments prompted us to study their metabolic niches in the honey bee gut environment. We carried out 16S rRNA gene-based community analysis, sequenced 12 isolates of Alpha2.1 and Alpha2.2, and carried out comparative genomic analysis including previously sequenced strains. We find that the two phylotypes are differentially distributed along the worker and queen bee gut, and confirm previous studies that show that they belong to distinct phylogenetic lineages within the Acetobacteraceae. The comparative genome analysis suggests that the two phylotypes have different strategies to metabolize carbon sources and to harvest energy. Moreover, we identified several characteristics of Alpha2.1 that are unique among Acetobacteraceae, which is in agreement with its position on a deep-branching lineage within the Acetobacteraceae.

## Materials and Methods

### Bee sampling

Four queen bees and four worker bees of the European honey bee *(Apis mellifera),* were sampled in Summer 2017. Three of each came from different colonies at the University of Lausanne and one from a professional beekeeper in Western Switzerland (Imkerei Giger). The guts were dissected into gut compartments (honey crop, midgut, ileum, and rectum), and homogenized as described in (Ellegaard and Engel 2019). Each sample was split in two, one of which was used for amplicon sequencing and the other for bacterial isolation. For the samples used for bacterial isolation, the four different gut regions were pooled together prior to plating them on different media.

### Amplicon sequencing

DNA from the different gut samples was isolated using an established CTAB/phenol extraction protocol (Kesnerova, et al. 2017). The 16S rRNA gene was amplified with primers 27F and 907R prior to sending the samples for amplicon sequencing analysis at Microsynth (Switzerland). At Microsynth, the V4 region of 16SrRNA gene was amplified using universal primers 515F/806R (Caporaso, et al. 2011), and the amplified fragments were purified and sequenced with the Illumina MiSeq platform (2×250bp). The number of 16S rRNA gene copies per host actin copy was quantified through qPCR with universal bacterial primers for honey bee gut (F 5’AGGATTAGATACCCTGGTAGTCC-3’, R 5’-YCGTACTCCCCAGGCGG-3’) following the method described by Kešnerová et al. (2017).

### Sequence processing and community analysis

Raw reads were processed and reads with more than 75% of bases below a quality score of 33 were filtered with FastX-Toolkit (Gordon & Hannon 2010, unpublished, http://hannonlab.cshl.edu/fastx_toolkit). Remaining paired reads were merged with PEAR (Zhang, et al. 2014). Paired reads were quality-filtered, dereplicated, clustered into OTUs at 97% identity and chimera-filtered with VSEARCH (Rognes, et al. 2016). OTU abundance was calculated by mapping the total quality-filtered paired reads to the final clusters using VSEARCH *--usearch_global*. The resulting abundances were normalized by the total 16S rRNA gene copy numbers as estimated through qPCR.

Representative OTU sequences were assigned to taxonomic categories with SINA (v.1.2.11) against SILVA_132_NR99 (Pruesse, et al. 2012), and composition was visualized with PHINCH v.1 (Bik 2014). Differences in copy numbers across samples were evaluated with an aligned Rank Transformation of a Factorial Model with the R package ARTool (Kay and Wobbrock 2016). Comparative abundances of Alpha2.1 and Alpha2.2 were expressed as log-ratios of the normalized copy numbers for all OTUs assigned to Alpha2.1 (*Commensalibacter*) and Alpha2.2 (*Bombella*).

### Bacterial culturing, DNA isolation, and genome sequencing

Serial dilutions of the gut homogenates were plated on Sabouraud Dextrose Agar (SDA) or MRS + Mannitol agar and incubated at 35°C in 5% CO2 incubator. After 3-5 days of incubation, single colonies were picked, restreaked on fresh agar, and incubated for another 2-3 days. Isolates were genotyped with universal bacterial primers and *rpoB*-specific primers (0937: GAAATTTATGCCGAGGCTGG; 0938: GAAATTTATGCCGAGGCTGG) as described in Ellegaard et al (2019), and stocked in MRS broth containing 25% glycerol at −80°C. A total of eight strains of Alpha2.1 and Alpha2.2 were selected for genome sequencing. Four additional strains were selected from a previous culturing effort using a similar culturing approach (see **Table S1**).

### Genome sequencing, assembly, and annotation

For Illumina genome sequencing, genomic DNA was isolated from fresh bacterial cultures using the GenElute Bacterial Genomic DNA Kit (SIGMA) according to manufacturer’s instructions. Genome sequencing libraries were prepared with the TruSeq DNA kit and sequenced on the MiSeq platform (Illumina) using the paired-end 2×250-bp protocol at the Genomic Technology facility (GTF) of the University of Lausanne. The genome sequence analysis was carried out as described in Ellegaard et al. (2019). In short, the resulting sequence reads were quality-trimmed with trimmomatic v0.33 (Bolger, et al. 2014) and assembled with SPAdes v.3.7.1 (Bankevich, et al. 2012). Small contigs (less than 500 bp) and contigs with low kmer coverage (less than 5) were removed from the assemblies, resulting in 5-22 contigs per assembly.

Two strains, ESL0284 and ESL0368, one from Alpha2.1 and Alpha2.2 each, were selected for sequencing with PacBio 20K (Pacific Biosciences) single-molecule realtime (SMRT) technology. High-molecular weight genomic DNA was extracted with the previously established CTAB/phenol extraction protocol (Kesnerova, et al. 2017). De novo genome assembly was done using the Hierarchical Genome Assembly Process (HGAP) version 2.3 (Chin, et al. 2013). These two completely assembled genomes served as references to order the contigs of the Illumina assemblies using MAUVE v2.4 (Rissman, et al. 2009). The origin of replication was determined based on the GC-skew and set to position 1 by cutting the contig at the corresponding location. The same was done for the published complete genome of Alpah2.1 strain AMU001 (Siozios, et al. 2019), but not for any of the other published draft genomes that were included in the analysis of the gene content (Corby-Harris and Anderson 2018; Smith and Newton 2018). Assembly quality was checked by remapping reads to assemblies with the Burrows-Wheeler Aligner (Durbin 2014). The genomes were annotated using the ‘Integrated Microbial Genomes and Microbiomes’ (IMG/mer) system (Chen, et al. 2017).

### Inference of gene families and genome-wide phylogeny

For the gene content analysis and the inference of the genome-wide phylogeny, we determined gene families, i.e. sets of homologous genes, across 56 genomes from the family Acetobacteraceae and other Alphaproteobactera using OrthoMCL (Li, et al. 2003). We included the 12 newly sequenced strains of Alpha2.1 and Alpha2.2, and 44 previously sequenced strains (including 9 previously sequenced Alpha2.1 and Alpha2.2 strains and 35 genomes of other Acetobacteraceae and closely related Alphaproteobacteria). Protein sequences of all CDS of the 56 genomes were searched against each other using BLASTP. BLASTP hits with an e-value <10^-5^ and a relative alignment length of >50% of the length of the query and the hit CDS were retained.

The OrthoMCL analysis was carried out as recommended with the mcl program run with the parameters ‘--abc -I 1.5’. A total of 11,451 gene families were identified. The remaining CDS (14,453 CDS) were singletons, i.e. they had no detectable homolog in any other genome in our dataset.

A total of 361 single copy orthologs were extracted from the OrthoMCL output (i.e. gene families having exactly one representative in every genome in the analysis). The protein sequences of each of these gene families were aligned with mafft (Katoh and Standley 2013) and alignment columns represented by <50% of all sequences removed. The single gene family alignments were concatenated and used as the basis for inferring a core genome phylogeny using RAxML (Stamatakis 2014) with the PROTCATWAG model and 100 bootstrap replicates.

### Comparison of genome structure and divergence

The R package genoplotR was used to compare and visualize whole genome alignments (Guy, et al. 2010). Pairwise BlastN comparison files were generated using command line blast v2.2.31+ using a bit score cutoff of 100. To estimate sequence divergence between genomes, we calculated pairwise ANI with orthoani (Lee, et al. 2016) using the exectutable “OAT_cmd.jar” with the parameter “-method ani.”.

### Comparison of the functional gene content

To analyze differences in gene content between Alpha2.1 and Alpha2.2, we extracted all OrthoMCL gene families that contained a homolog in at least one of the 21 genomes of the two phylotypes of interest and included also singletons, i.e. gene families that had homologs in only one of the genomes. This resulted in a total of 3,275 gene families identified across 21 genomes of Alpha2.1 and Alpha2.2. These gene families were categorized into ‘shared’ and ‘specific’ core and pan genome subsets depending on their presence/absence in the genomes of Alpha2.1 and the two sub-lineages of Alpha2.2. To analyze functional differences between different subsets of core gene families, we assigned the IMG/mer annotation of one of the homologs (if possible the one of the complete reference genomes) to each gene family. These annotations were used to determine the distribution of gene families into COG categories and to identify the shared and phylotype-specific metabolic capabilities, biosynthetic pathways, and transport functions based on the analysis of KEGG pathways. For the analysis of the respiratory chain, we combined several approaches: The characterization of the electron acceptors was mainly based on KEGG annotations and the previous publication of the genome of *Bombella intestini* (Li, et al. 2016). To identify respiratory dehydrogenases we carried out keyword searches with ‘dehydrogenase’ and ‘reductase’, and identified all gene families that belonged to an enzyme class (EC number) that has been described in the literature as being a respiratory enzyme (Marreiros, et al. 2016). The TCA cycle analysis was based on KEGG annotations. For inferring phylogenetic trees of the nitrate reductase subunit alpha (*narG*) and the nitric oxide reductase, the amino acid sequence of one of the homologs was searched against the nr database using BLASTP. From the top blast hits, a subset of homologs identified in divergent strains and species was selected to build a phylogeny using RAXML. The most similar homologs present among Acetobacteraceae were identifiedby carrying out a second BLASTP search against nr by restricting hits to the family Acetobacteraceae (taxid:433).

## Results

### Community analysis of the gut microbiota of worker bees and queens suggests different niches of Alpha2.1 and Alpha2.2

Bacteria of the Acetobacteraceae Alpha2.1 and Alpha2.2 have been reported to be dominant community members of the gut microbiota of adult honey bee queens (Anderson, et al. 2018; Corby-Harris, et al. 2014; Kapheim, et al. 2015; Tarpy, et al. 2015), and can also be abundant in the crop of worker bees (Corby-Harris, et al. 2014). To corroborate these previous results and obtain additional insights about the distribution and relative contribution of these phylotypes to the gut microbiota, we determined total bacterial loads and community composition in four different gut compartments (crop, midgut, ileum, and rectum) of honey bee queens and worker bees.

qPCR analyses revealed significant differences in total 16S rRNA gene copy number across castes (i.e. queens and workers) and gut regions (*F_caste_*=15.7, *F_gut_*=13.9, *p*<0.05, **Figure 1A**), with the largest difference being that queens had significantly smaller bacterial loads than workers. The largest bacterial loads were found in the rectum, and the smallest in the crops, as reported in previous studies (Martinson, et al. 2012; Powell, et al. 2018). The only gut region where the queens had significantly larger loads than the workers was the crop (**Figure 1A**).

**Figure 1.**
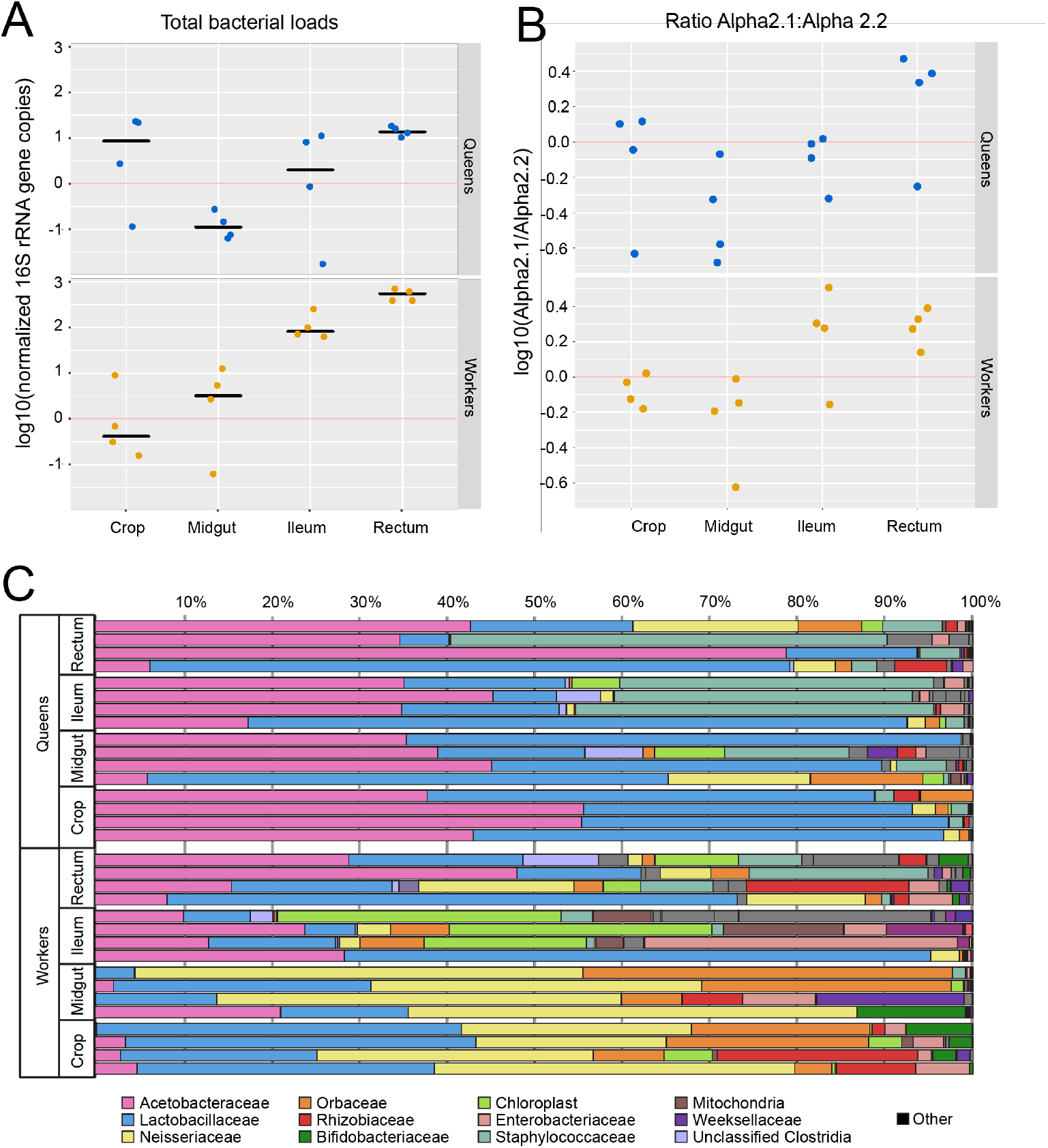
Community analysis of different gut compartments of adult worker and queen bees (n=4 for each category). (A) Total bacterial loads in different gut compartments assessed by qPCR using universal 16S rRNA gene primers and expressed as 16SrRNA copies per host actin copy. (B) Abundance of Alpha2.1 relative to Alpha2.2 in the four gut compartments of worker and queen bees. (C) Bar plot depicting the community composition per gut compartment and caste as relative abundance at the family level.

The community composition differed markedly between castes and gut regions. Queen guts were characterized by the dominance of family Acetobacteraceae (i.e. Alpha2.1 and Alpha2.2) across all compartments and a low abundance of *Snodgrassella (Neisseriaceae)* and *Gilliamella (Orbaceae),* corroborating previous studies (**Figure 1C**) (Anderson, et al. 2018; Corby-Harris, et al. 2014; Kapheim, et al. 2015; Tarpy, et al. 2015). The family Acetobacteraceae was also present in all gut compartments of worker bees, but was proportionally much more abundant in the crop and midgut compared to midgut and rectum. Taking the bacterial loads per compartment into consideration, the family Acetobacteraceae was most abundant, and contributed the most to total community composition, in the rectum of queens (**Figure 1A and 1C**).

None of the two Alpha2 phylotypes were exclusively associated with queen guts, but instead were found in all gut compartments from both castes. The average 16S rRNA gene copies per actin copy of both phylotypes were consistently higher in workers than in queens, and overall Alpha2.1 was higher than Alpha2.2. Nevertheless, we found differences in the ratios between Alpha2.2 to Alpha2.1 within gut compartments: Alpha2.1 was more abundant in the rectum, and Alpha 2.2 was more abundant in midguts of both queens and workers (**Figure 1B**). The ratios in the ileum differed between castes, with Alpha2.2 being more abundant in the queens, and Alpha2.1 being more abundant in the workers. As expected between individual bees, we observed large variability across the four replicates, particularly for queen’s crops, but the pattern observed in queens is also in line with that reported in Anderson et al. (2018).

In summary, both phylotypes are found across all gut compartments of queens and worker bees, but while Alpha2.2 seems to be more abundant and contributes proportionally more to the community in the queen crop and midgut, Alpha2.1 is much more abundant in the rectum of both castes.

### Alpha2.1 and Alpha2.2 belong to two distinct phylogenetic clades within the Acetobacteraceae

To facilitate functional and phylogenetic analysis, we sequenced the genomes of seven Alpha2.1 and five Alpha2.2 strains (**Table S1**). Eight of the 12 strains were isolated from the same worker and queen bees that were analyzed in the previous section. The other four strains were isolated from other worker and queen bees. One strain of each phylotype was assembled into a single circular chromosome (strain ESL0284 for Alpha2.1, and strain ESL0368 for Alpha2.2) and served as reference for our analysis. The assemblies of the other genomes consisted of 5-22 contigs. Genome size varied little among strains and was comparable between the two phylotypes ranging from 1.85-2.07 Mb. The overall genome structure was largely conserved in both phylotypes as based on whole genome alignments with the complete genome of the reference strain of each phylotype (**Figure S1**).

To better understand the phylogenetic relationship between Alpha2.1, Alpha2.2, and other Acetobacteraceae, we inferred a genome-wide tree of the 12 newly sequenced Alpha2.1 and Alpha2.2 strains, nine previously sequenced Alpha2.1 and Alpha2.2 strains (Corby-Harris and Anderson 2018; Siozios, et al. 2019), and 37 strains of other Alphaproteobacteria (mainly Acetobacteraceae). This tree revealed that Alpha2.1 and Alpha2.2 are polyphyletic, i.e. belong two distinct clades within the Acetobacteraceae (**Figure 2**).

**Figure 2.**
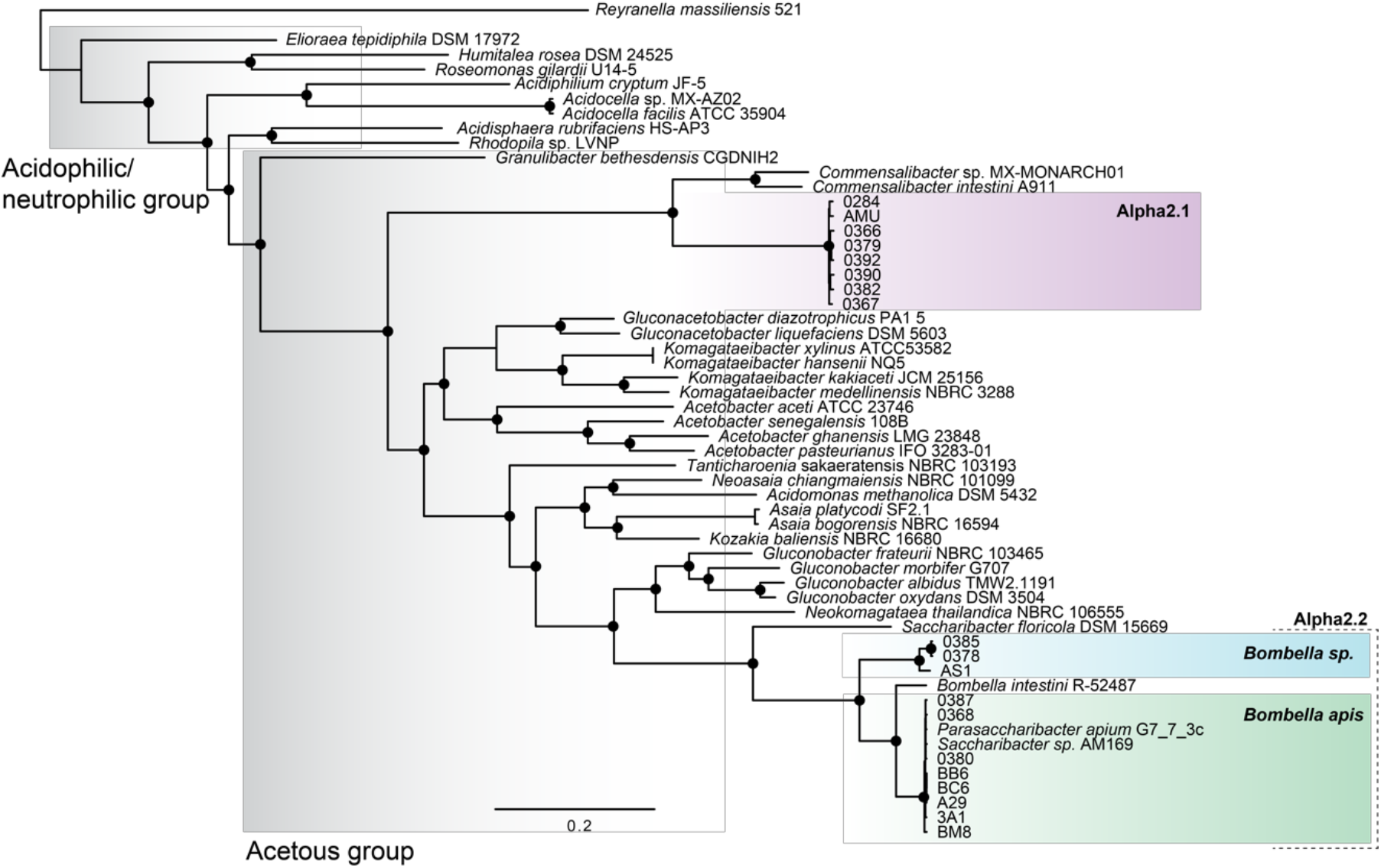
Genome-wide phylogeny of Acetobacteraceae based on 361 single-copy core genes. The tree was inferred using maximum likelihood. Filled circles indicate bootstrap value of 100 out of 100 replicates. Branches without circles indicate bootstrap values <80. The length of the bar indicates 0.2 amino acid substitutions per site. The three clades of interest are highlighted in magenta, blue, and green. The acetous and acidophilic/neutrophilic groups are indicated by shades of grey. Accession numbers of the Alpha2.1 and Alpha2.2 strains can be found in Table S1.

Alpha2.1 formed, together with the two previously identified strains of the genus *Commensalibacter,* a deep-branching monophyletic lineage, which was basal to most AABs within the Acetobacteraceae. However, a relatively long branch separated Alpha2.1 from the two *Commensalibacter* strains (**Figure 2**) and the average nucleotide identity (ANI) between their genomes was very low (~70%, see **Table S2 and Figure S2**) indicating deep divergence. In contrast, the analyzed strains of Alpha2.1 were closely related to each other, as evident from the short branches separating the different strains in the phylogenetic tree (**Figure 2**) and the high ANI between their genomes (98-99%, see **Table S2 and Figure S2**). This agrees with a recent metagenomic study, which found that Alpha2.1 belongs to the species with the lowest extent of strain-level diversity within and between individual honey bees (Ellegaard and Engel 2019). Our results suggest that Alpha2.1 presents a novel species of the genus *Commensalibacter*. Notably, the genomes of Alpha2.1 and the two strains of *Commensalibacter* had a very low GC content (~37%) relative to all other sequenced Acetobacteraceae, which is usually between 50-60% (**Table S1**), providing further evidence for their distinctive position within this family.

In contrast to the basal position of Alpha2.1 in the acetous group of the Acetobacteraceae, our phylogeny revealed that Alpha2.2 is nested within the *Gluconobacter* clade and is most closely related to *S. floricola* DSM 15669. The sequenced Alpha2.2 strains clustered into two sub-lineages (**Figure 2**) that were separated by a well-supported branch leading to *Bombella intestini* R-52487, a species previously isolated from the gut of a bumble bee (**Figure 2**) } (Li, et al. 2015). ANI values between strains of the same sub-lineage were relatively high (92-99%), while ANI values between strains of the two different sub-lineages were low (74-75%, see **Table S2 and Figure S2**). No genome sequence is currently available for the designated type strain MRM of the described species *B. apis* (Yun, et al. 2017). However, based on 16S rRNA sequence similarity, this strain could undoubtedly be assigned to one of the two sub-lineages (**Figure S3**). We will refer to this sub-lineage as ‘*Bombella apis*’, while sub-lineage A2.2-2 may represent a new species of the genus *Bombella*.

Taken together, we can conclude that while the sequenced *Bombella* strains clearly belong to the acetous group, more precisely to the *Gluconobacter* clade, the deep-branching basal position of Alpha2.1 suggests that it belongs either to an early diverging clade within the acetous group, or to a yet unexplored third subgroup. This calls for further taxonomic analyses, since the atypical GC content of the Apha2.1 genomes might confound the position in the phylogeny.

### Functional similarities between Alpha2.1 and Alpha2.2 based on the shared core gene content

To compare the functional gene content of Alpha2.1 and Alpha2.2, we clustered homologous genes of the two phylotypes into gene families and looked at their distribution across the 21 analyzed genomes. A total of 3,275 gene families were identified (including singletons), of which 1,006 were conserved across all strains, hereafter referred to as the shared core genome of Alpha2.1 and Alpha2.2 (**Figure 3A, Table S3**).

**Figure 3.**
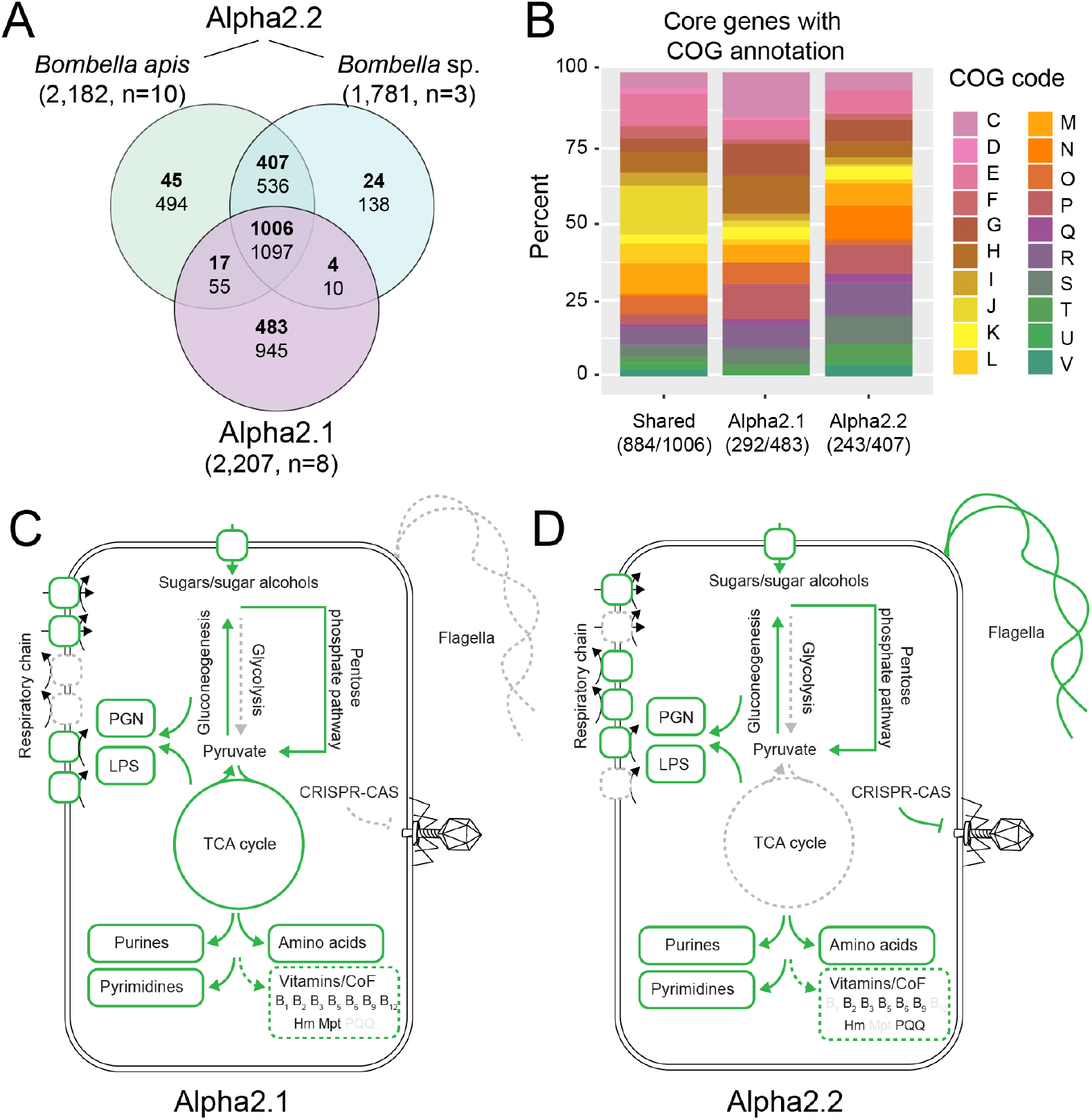
Distribution of gene families among the phylotypes Alpha2.1 and Alpha2.2 and overview of their major functional capabilities. (A) Venn diagram depicting shared and unique gene families in the three phylotypes (Alpha2.1 and the two Alpha2.2 sublineages *Bombella apis* and *Bombella sp.).* Gene families present in all genomes of a given group (i.e. core genes) are indicated in bold font, and gene families present in at least one genome (i.e. pangenome) are indicated in normal font. Numbers in brackets indicate total gene families found for each clade. Notably, the number of analyzed genomes (indicated by n) can influence the number of pan and core gene families. (B) Distribution of shared and phylotype-specific core gene families grouped by COG categories. Numbers in brackets indicate number of core genes with annotation and total number of core genes. (C) and (D) Summary of the major functional capabilities of Alpha2.1 and Alpha2.2 based on the shared and the phylotype-specific gene content. Dashed lines indicated pathways and biosynthetic capabilities missing in the respective phylotype. Solid lines indicate the presence of pathway. LPS, lipopolysaccharide, PGN, peptidoglycan, CoF, cofactor; Hm, heme; MPT, molybdopterin; PQQ, pyrroloquinoline quinone.

Inspection of the shared core genome revealed that Alpha2.1 and Alpha2.2 have similar biosynthetic capabilities as other proteobacterial bee gut symbionts (Kwong and Moran 2016). Both phylotypes encode complete gene sets for the biosynthesis of all amino acids (except for alanine), peptidoglycan, LPS, and five vitamins (riboflavin (vitamin B_2_), nicotinate (vitamin B_3_), pantothenate (vitamin B_5_), pyridoxine (vitamin B_6_), and tetrahydrofolate (vitamin B_9_)) (**Figure 3C** and **3D**, **Figure S4**). In contrast to most symbionts in the honey bee gut, Alpha2 lineages code for pathways for the *de novo* synthesis of purine and pyrimidine nucleosides. The dihydroorotate dehydrogenase (E.C. 1.2.5.2) in Alpha2.1 is quite divergent, with higher identity to its homologues in *Gilliamella* (~64%) than those in Alpha2.1 (~24%). The gene is adjacent to *uup*, involved in transposon excision in *E. coli* (Carlier, et al. 2012), suggesting it has been acquired by HGT. Notwithstanding, all genomes from Alpha2.1 code for a symporter for the uptake of environmental orotate, suggesting it can also satisfy its need for this nucleoside precursor extracellularly. The *de novo* nucleoside biosynthesis is costly, and several gut symbionts lack these functional pathways, preferring their uptake from the environment as evidenced by nucleoside depletion in the bee gut in the presence of microbiota (Kesnerova, et al. 2017). Alpha2.2 genomes code for an adenosine importer, which in turn can be interconverted with inosine, xanthine and guanosine. This suggests that Alpha2.2, like most other bee gut symbionts, prefers to acquire these DNA and RNA building blocks from the environment rather than synthesizing them *de novo*.

All analyzed strains encoded a complete gene set of the Embden-Meyerhof-Parnas (EMP) pathway except for a homolog of the 6-phosphofructokinase gene (*pfkA*, EC:2.7.1.11), indicating that Alpha2.1 and Alpha2.2 are both capable of performing gluconeogenesis but incapable of carrying out glycolysis. However, all strains harbored the necessary genes to incorporate sugars and sugar acids via the Pentose phosphate pathway (PPP) into intermediate cellular metabolites (**Table S4, Figure 3C** and **3D**, **Figure S4**). In addition, all strains of Alpha2.2 carried a 6-phosphogluconate dehydratase (*edd*, EC 4.2.1.12) and a 2-keto-3-deoxy-6-phosphogluconate (*eda*, EC 4.1.2.14), which allows them to use the Entner-Doudoroff pathway (ED).

Analysis of the shared core genome content also revealed that both phylotypes - like most other Acetobacteraceae (Matsushita and Matsutani 2016)-carry out oxidative phosphorylation and rely on oxygen as the final electron acceptor (**Figure S4, Figure 4A, Table S4**). They encoded genes of the respiratory complex III ubiquinol:cytochrome *bc1* reductase, which can transfer electrons from the quinone pool to different cytochrome *c*-containing proteins. While they lacked genes for the complex IV COX-type cytochrome c terminal oxidase (*coxABC*), both lineages carried genes coding for two ubiquinol-dependent terminal oxidases: *ctaAB* and *cyaDCBA* for the cytochrome *bo3* oxidase, and *cydAB* for the low oxygen affinity cytochrome *bd* oxidase (Matsushita and Matsutani 2016). Overall, this suggest that both lineages heavily rely on utilizing the ubiquinone pool to transfer electrons.

**Figure 4.**
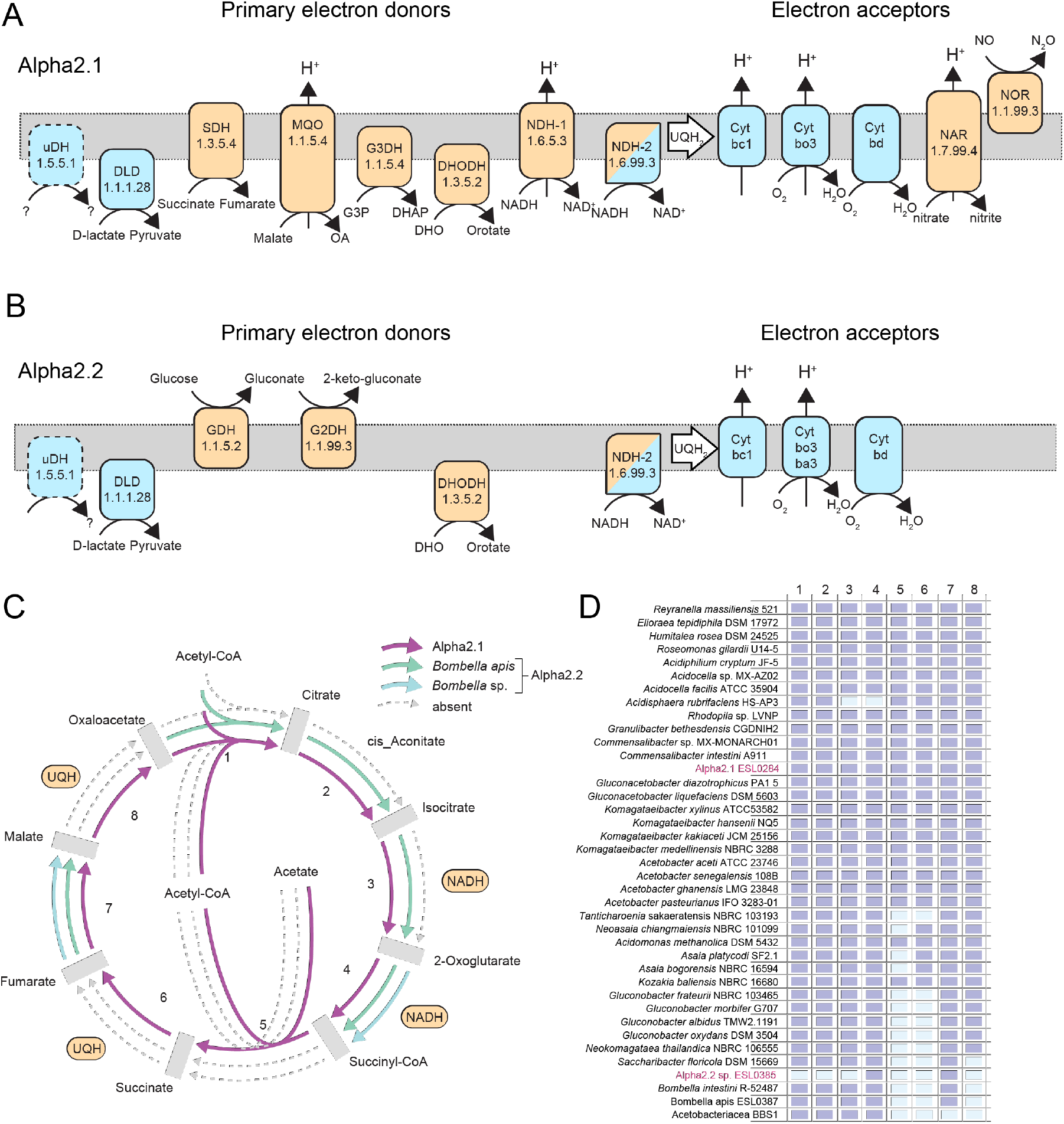
Respiratory chain of Alpha2.1 (A) and Alpha2.2 (B). Shared components are shown in blue, phylotype-specific functions are shown in tan. Dashed outlines indicate that this function is not conserved in all sequenced strains. Both colors indicate the presence of two copies of this function, one which is phylotype-specific, another one which is shared among the two. UQH2 depicts the ubiquinone pool (C) TCA cycle completeness in Alpha2.1 and both Alpha2.2 sub-lineages, *Bombella apis* and *Bombella* sp. Steps where electrons are transferred to ubiquinone (UQH) and NADH are are indicated. (D) Completeness of the TCA cycle across Acetobacteraceae strains shown in Figure 1, including a recently described strain isolated from the gut of an ant, Acetobacteraceae BBS1, which also has an incomplete TCA cycle (Brown and Wernegreen 2019). DLD, D-lactate dehydrogenase; SDH, succinate dehydrogenase; MQH, maltate:quinone oxidoreductase; G3DH, glycerol-3-phosphate dehydrogenase; DHOHD, dihydroorotate dehydrogenase; NDH-1, NADH dehydrogenase type 1; NDH-2, NADH dehydrogenase type 2; NAR, nitrate reductase; NOR, nitric oxide reductase; uDH, unknown flavoprotein dehydrogenase; GDH, PQQ-dependent glucose dehydrogenase; G2DH, gluconate 2-dehydrogenase, UBQ_2_. ubiquinone-2; Cyt, cytochrome. 1, Citrate synthase; 2, Aconitate hydratase; 3, Isocitrate dehydrogenase; 4, 2-oxoglutarate dehydrogenase; 5, Succinyl-CoA:acetate CoA-transferase (*aarC*); 6, Succinate dehydrogenase; 7, Fumarase class II; 8, Malate dehydrogease (*mqo*).

One of the hallmarks of the Acetobacteraceae metabolism is the production of acetate from ethanol via acetaldehyde (De Roos and De Vuyst 2018). Although both lineages carried genes for NADH-producing alcohol dehydrogenases to convert ethanol into acetaldehyde (EC:1.1.1.1, EC:1.1.1.284,) none of them was a homolog of the ubiquinone-dependent PQQ alcohol dehydrogenases characteristic of acetate production and responsible in *Acetobacter pomorum* to induce the host insulin signaling pathway of *Drosophila* resulting in increased growth and development (Shin, et al. 2011). Moreover, none of the lineages carried genes for the subsequent production of acetaldehyde to acetate (EC:1.2.1.-), nor for acetate transport (COG1584). This means that none of the lineages produce ethanol through oxidative fermentation nor are they acetate producers, two characteristic reactions of Acetobacteraceae.

### Functional differences between Alpha2.1 and Alpha2.2 based on the phylotype-specific gene content

Although sharing many metabolic capabilities, Alpha2.1 and Alpha 2.2 each harbored a considerable number of phylotype-specific core gene families, i.e. gene families present across all analyzed strains of one phylotype, but absent from all strains of the other phylotype (**Figure 3A, Table S5**): 483 and 407 gene families for Alpha2.1 and Alpha2.2, respectively. The number of core gene families specific to each of the two sub-lineages of Alpha2.2 was relatively small: 45 gene families for *B. apis* and 24 gene families for *Bombella sp.* All three groups (i.e. Alpha2.1, *B. apis,* and *B. sp.)* also harbored a relatively large flexible gene pool (i.e. gene families present in only asubset of the strains), as indicated by the total number of gene families in each group (i.e. pan genome) as compared to the core genome (**Figure 3A, Table S5**).

The 483 and 407 phylotype-specific core gene families belonged to a wide range of COG categories suggesting differences in diverse metabolic functions. However, four COG categories stood out as being particularly abundant among the Alpha2.1-specific core gene content as compared to the shared core gene content (**Figure 3B**): ‘Energy production and conversion’ (COG C), ‘Carbohydrate transport and metabolism’ (COG G), ‘Coenzyme transport and metabolism’ (COG H), and ‘Inorganic ion transport and metabolism’ (COG P). Gene families of these particular categories encoded a relatively large number of dehydrogenases/oxidoreductases, putative transporters for nitrate/nitrite, sulfate, and iron, and nearly complete gene sets for the synthesis of vitamin B_1_ (thiamine), vitamin B12, and the co-factor molybdopterin, all of which were absent from Alpha2.2. Interestingly, several genes linked to the TCA cycle were also specific to Alpha2.1 (**Figure 3C and 3D**).

Among the Alpha2.2-specific core gene content two COG categories were particularly abundant compared to the shared core genome (**Figure 3B**): ‘Inorganic ion transport and metabolism’ (i.e. COG P) and ‘Cell motility’ (i.e. COG N). Almost all gene families in the category ‘Cell motility’ coded for different flagella subunits, suggesting that Alpha2.2, but not Alpha2.1, is motile (**Figure 3C and 3D**). Gene families in ‘Inorganic ion transport and metabolism’ encoded for diverse transporters, in particular for iron and phosphate. All *Bombella* strains also harbored genes encoding the redox cofactor pyrroloquinoline quinone (PQQ) and a PQQ-dependent glucose dehydrogenase involved in respiration (see below). Another notable difference between the two phylotypes was that Alpha2.2 encoded an entire CRISPR/CAS9 system, while Alpha2.1 was lacking any homolog of these antiviral defense systems (**Figure 3C and 3D**).

### Major differences in respiratory chain and TCA cycle between Alpha2.1 and Alpha2.2

Both phylotypes seem to carry out aerobic respiration to gain energy. However, the presence of various dehydrogenases and TCA cycle genes among the phylotype-specific gene content suggested differences in the energy metabolism (**Table S5**). We identified 16 membrane-associated dehydrogenase/reductases likely to be involved in electron transport respiratory chain hence the production of energy (**Figure 4**, **Table S6**).

Six dehydrogenase/reductase gene families were present among the shared gene content of Alpha2.1 and Alpha2.2, three of which are electron donors (a D-lactate dehydrogenase, a complex I type-II NADH dehydrogenase, and a putative membranebound dehydrogenase), and three of which are electron acceptors, namely the ubiquinol:cytochrome *bc1* complex III and the two terminal electron acceptors cytochrome *bo3* and *bd* ubiquinol oxidases. and *bo3*). These consist of the common respiratory metabolism shared between the two lineages. The remaining 11 dehydrogenases/reductases belonged to the phylotype-specific core gene content (i.e. were present in all strains of one phylotype but absent from the other).

Eight of these phylotype-specific dehydrogenases/reductases were only present in Alpha2.1. Six dehydrogenases are electron donors to ubiquinone from oxidation of succinate, NADH (one *nuo/*type-I in addition to the shared *ndh*/type-II), glycerol-3-phosphate, malate and the aforementioned dihydroorotate dehydrogenase(**Figure 4A**, **Table S6**). The type I NADH dehydrogenase and the nitrate reductase are proton-pumping enzymes that directly contribute to the production of energy (Marreiros, et al. 2016). Both enzyme complexes are dependent on the cofactor molybdopterin, which explains the presence of the corresponding biosynthesis genes in the genomes of Alpha2.1, but not Alpha2.2. The genes encoding the co-factor and the nitrate reductase are located in the same genomic island and have best blast hits to Gamma- and Betaproteobacteria, suggesting acquisition by HGT (**Figure S5**). (**Figure 4A**).. The other two terminal oxidases are nitrate and nitric oxide reductases, which suggest that Alpha2.1 has the capability to carry out anaerobic respiration (Figure 4A). Homologs of the gene encoding the nitric oxide reductase were not present in any other closely related *Acetobacteriaeceae* strain (i.e. other *Commensalibacter* sp.) suggesting a specific role for Alpha2.1 and acquisition by HGT as is common for denitrification enzymes (Jones, et al. 2008) (**Figure S6**).

Only three dehydrogenases were specific to Alpha2.2: a different dihydroorotate dehydrogenase, the above mentioned PQQ-dependent glucose dehydrogenase, and a gluconate-2-dehydrogenase (**Figure 4B**). The last two catalyze the periplasmic conversion of glucose into gluconate and gluconate into 2-keto-gluconate, and are characteristic of the oxidative fermentation pathway common to most Acetobacteraceae. This is in contrast to Alpha2.1, which lacked genes encoding periplasmic dehydrogenases and hence seems not able to carry out oxidative fermentation.

We also identified major differences in the TCA cycle between the two phylotypes. All strains of Alpha2.1 harbored the full gene set of the TCA cycle, including an acetate:succinate CoA-transferase gene (*aarC*) for metabolizing acetate to acetyl-CoA (Mullins, et al. 2008), a typical feature of the TCA cycle of Acetobacteraceae thought to be an adaptation to the high amounts of acetate produced by some of them (**Figure 4C**). In contrast, the TCA cycle was incomplete across strains of Alpha2.2 to different degrees. While in the sub-lineage of *B. apis* the enzymatic steps from succinyl-CoA to fumarate (two steps) and from malate to oxaloacetate (one step) were missing, the strains of the other sublineage were also missing genes for the conversion of acetyl-CoA to 2-oxoglutarate (four steps). Several Acetobacteraceae have been reported to harbor partial or modified TCA cycles, especially those that utilize oxidative fermentation to gain energy (Brown and Wernegreen 2019; Mullins, et al. 2008). However, in none of them the pathway seems to be as reduced as in this particular sublineage of *Bombella* (**Figure 4D**).

Intriguingly, the fate of malate in the two phylotypes is markedly different. Alpha2.1 displays a malate:quinone oxidoreductase as part of the conversion of malate to oxaloacetate in the TCA cycle. This enzyme is an alternative that contributes to both the proton motive force through proton pumping and the ubiquinone pool with electrons. Malate can also be converted into pyruvate and CO2 through malate dehydrogenases. Alpha2.1 codes for the NADP-dependent *maeB* (EC 1.1.1.40), and Alpha2.2 codes for a NAD-dependent *maeA* (EC 1.1.1.38). The latter is found adjacent to the class-II fumarase *fumC* (EC 4.2.1.2) in the Alpha2.2 genomes, that also lack the genes to generate fumarate from succinate, and hence must rely on fumarate produced through other anabolic pathways such as the urea cycle or aspartate metabolism.

## Discussion

The two Acetobacteraceae, Alpha2.1 and Alpha2.2, are common members of the bee gut microbiota. They have been repeatedly identified by 16S rRNA gene sequence analyses in samples from the bee gut or the hive environment and several studies have provided important insights about their ecology (Cox-Foster, et al. 2007; Kapheim, et al. 2015; Kešnerová, et al. 2019; Kwong, et al. 2017; Martinson, et al. 2011; Martinson, et al. 2012; Moran, et al. 2012; Powell, et al. 2018; Vojvodic, et al. 2013). However, a comparative analysis between their genomes had not been carried out. Our study provides new insights about the phylogenetic relationship of Alpha2.1 and Alpha2.2, their functional gene content, and their metabolic and genomic differences.

We show that Alpha2.1 and Alpha2.2 belong to two distant phylogenetic clades within the Acetobacteraceae, confirming previous findings (Martinson, et al. 2011), and determining their taxonomic position more precisely. Alpha2.2 is part of the acetous group and falls within the genus *Bombella*, to which also *B. intestini* belongs (Li, et al. 2015). However, the Alpha2.2 strains isolated from honey bees belong to two divergent sub-lineages within the genus *Bombella.* The species formally described as *B. apis* (Yun, et al. 2017) and strains referred to as *Candidatus* ‘Parasaccharibacter apium’ and the undescribed strains *Saccharibacter* sp. 3.A.1 (Veress, et al. 2017) belong to the same sub-lineage, are identical or nearly identical in the 16S rRNA gene, and share high ANI with each other. While we acknowledge that the name ‘Parasaccharibacter apium’ has been proposed first (Corby-Harris, et al. 2014), it has never been formally described as a novel genus, and thus we propose that these closely related strains should be consistently referred to as *Bombella apis* (Yun et al. 2017).

The second sub-lineage of Alpha2.2 represents a putative novel species within the genus *Bombella* based on its genomic divergence and paraphyletic position relative to *B. apis*. The TCA cycle is almost completely lost in this sub-lineage suggesting distinctive functional capabilities as compared to *B. apis*. How prevalent this sublineage is across honey bee colonies, and whether it colonizes a different niche than *Bombella apis* remains to be determined.

The second phylotype, Alpha2.1, belongs to a highly divergent lineage within the Acetobacteraceae and is likely to represent a novel species within the genus *Commensalibacter*. The only three taxa which are known from this lineage have been isolated from insect guts, and all three display an atypical GC content compared to other Acetobacteraceae. Future sampling of bacteria from this lineage will help to solidify its position within the Acetobacteraceae phylogeny and to test if this lineage harbors exclusively insect gut-associated symbionts.

Despite the fact that Alpha2.1 and Alpha2.2 are both Acetobacteraceae and can colonize the same environment (i.e. the bee gut), we found major differences in their metabolic and respiratory capabilities. While both rely on the ubiquinone pool and oxygen- and ubiquinone-dependent terminal oxidases (and hence are aerobic), Alpha2.1 is also able to respire nitrate and nitric oxide, making it a facultative anaerobe.

The most important metabolic difference between the two phylotypes lies in their energy and central carbon metabolism. Alpha2.2 exhibits a simple metabolism that relies on the periplasmic oxidative fermentation of glucose and gluconate. Although it carries genes to incorporate these intermediates through the ED or PP pathways, this is unlikely, since most Acetobacteraceae accumulate 5-keto-D-gluconate extracellularly (Bringer and Bott 2016), including *B. apis* (Yun, et al. 2017), and it has been shown that a very small proportion of glucose and gluconate is incorporated into the cell during oxidative fermentation in *Gluconobacter oxidans*.,. This, together with a highly reduced TCA cycle that appears to retain only steps involved in other biosynthetic pathways, suggest that Alpha2.2 has a streamlined and rigid metabolism optimized for low-yield but rapid energy production from glucose without incorporation into biomass. This has been shown to occur when growth rate is slower than the reducing equivalent production rate in marine cyanobacteria (Braakman, et al. 2017), and may provide Alpha2.2 strains an advantage in glucose-rich environments, such as royal jelly (Simo and Christensen 1962). Worker bees secrete glucose oxidase into royal jelly to partly convert glucose into gluconolactone and hydrogen peroxide for antimicrobial purposes (Fratini, et al. 2016; Ohashi, et al. 1999). Alpha2.2 is able to further oxidize gluconolactone into gluconate (Smith and Newton 2018) and enter the ED pathway, potentially facilitating its growth in royal jelly.

In contrast to Alpha2.2, Alpha2.1 displays a wider range of primary electron donors that includes several organic acids and NADH, but does not permit oxidative fermentation. It carries the complete acetate-driven TCA cycle typically found inAcetobacteraceae, which is a low-yield variant already optimized for rapid resource consumption and replenishment of TCA intermediates while retaining steps needed to provide precursors for multiple biosynthetic pathways. A larger range of carbon substrate utilization fits a lifestyle that is more specialized to the honey bee hindgut. In support of this, a recent study showed that while many transient bacteria (including Alpha2.2) diminished or completely disappeared in long-lived winter bees, Alpha2.1 persisted in the gut environment and even significantly increased relative to foragers and nurses sampled in summer (Kešnerová, et al. 2019). The same pattern was also observed for another community member of the bee gut microbiota, *Bartonella apis.* Like Alpha2.1 this species relies on aerobic respiration (Segers, et al. 2017),in contrast to most other members, which are anaerobic fermenters. However, the metabolic or physicochemical conditions in the hindgut of winter bees that favor the growth of Alpha2.1 or *Bartonella apis* over other microbiota members is currently unclear.

In summary, we have shown that the two honey bee-associated Acetobacteraceae phylotypes both display adaptations towards fast growing metabolism, but in two markedly different ways. Alpha2.1 harvests energy from a broad-range of substrates and links substrate utilization with a flexible metabolism of oxidative and biosynthetic pathways, whereas Alpha2.2 has streamlined its metabolism by using oxidative fermentation for rapid energy harvesting from glucose, almost exclusively because of the loss of alternative oxidative pathways. Our results exemplify how physiological and metabolic differences may drive niche differentiation in terms of resource utilization and spatial distribution, facilitating the coexistence of phylogenetically related bacteria in the honey bee gut.

## Acknowledgments

Participants of the Master in Molecular Life Sciences at the University of Lausanne, class 2017-2018 (Marie-Pierre Meurville, Maxime Brunner, Jonathan Klopfenstein, Baptiste Micheli, Tess Bonato, Alice Wallef, Alicia Zufferey, Andrea Dos Santos, Augustine Jaccard, Béatrice Tappy, Boris Schnider, Christèle Aubry, Clara Heiman, Dinis Barros, Emilie Ha, Giti Ghazi Soltani, Henri Paratte, Ilaria Bernabei, Imre Banlaki, Julien Dénéréaz, Karim Saied, Liza Darrous, Lucas Beurret, Lucas Serra Moncadas, Lucien Roesch, Ludivine Brandt, Lydia Horwath, Margot Delavy, Maria Chadiarakou, Maria Hadres, Mathieu Quinodoz, Maxence Trottet, Pascal Roth, Perrine Steffe, Robin Hofmeister, Béatrice Tappy, and Vishwachi Tripathi) conducted preliminary research as part of the ‘Sequence-a-genome’ (SAGE) graduate course. We thank the School of Biology of the University of Lausanne for financial support, and the Competence Centre in Bioinformatics and Computational Biology of the Swiss Institute of Bioinformatics (SIB) for providing access to high-performance computation through the Vital-IT cluster (http:vital-it.ch). This research was funded through the School of Biology of the University of Lausanne, and the European Research Council (ERC-StG ‘MicroBeeOme’), the Swiss National Science Foundation (grant number 31003A_160345 and 31003A_179487) and HFSP (HFSP Young Investigator grant RGY0077/2016) received by PE.

## Data deposition

Genome sequences and Illumina data underlying genome assemblies and 16S rRNA gene amplicon analysis have been deposited on NCBI under Bioproject accession PRJNA589199. Codes and data of the analysis can be found via the following SWITCHdrive link: https://drive.switch.ch/index.php/s/syuWZAQKKGmQn0h. These files will be moved to a public Zenodo repository upon publication.

