## Supplementary Figures for "Acetobacteraceae in the honey bee gut comprise two distant clades with diverging metabolism and ecological niches"

### Supplementary Material

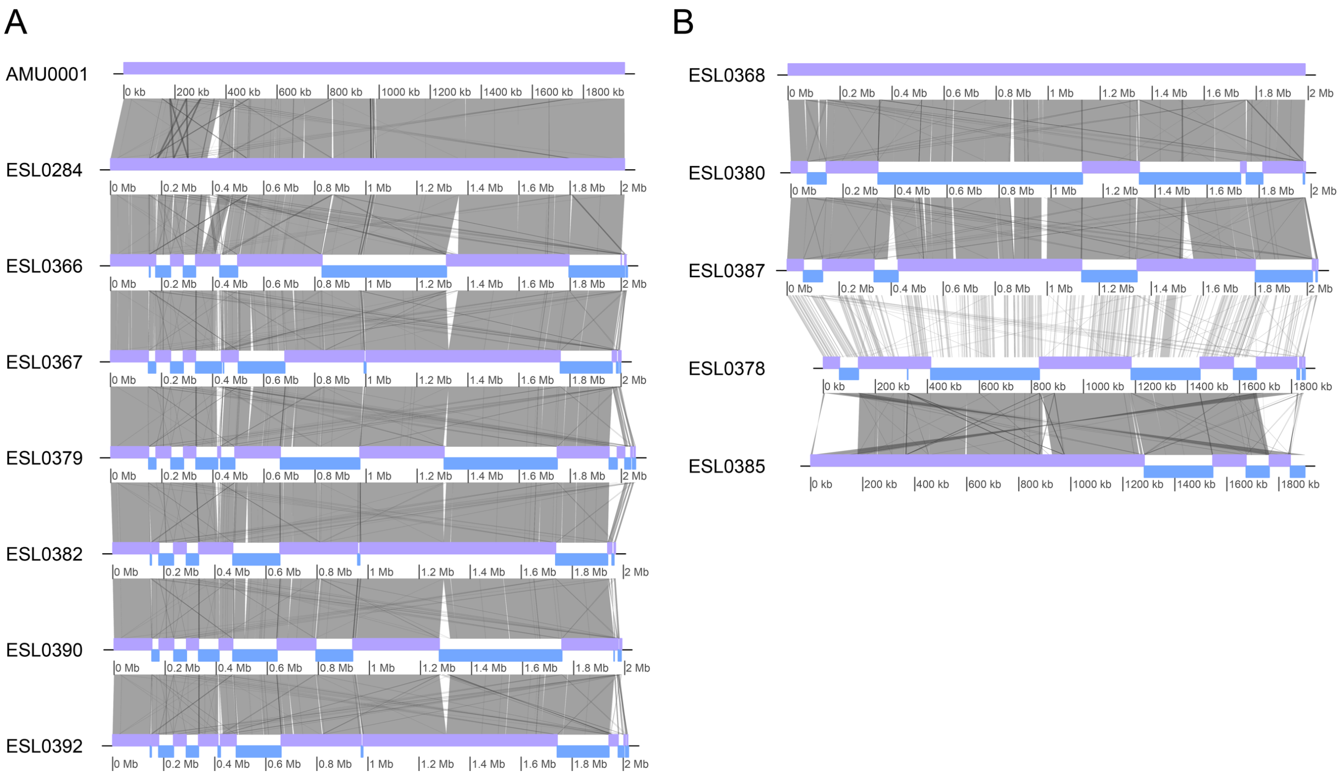


**Figure S1. Whole genome alignments of Alpha2.1 (A) and Alpha2.2 (B) genomes sequenced in this study.** Grey vertical lines present BlastN results with an e-value cutoff of $10^{-15}$, except for the comparison of ESL0387 and ELS0378, which belong to distinct sub-lineages of Alpha2.2 and for which an e-value cutoff of $10^{-5}$ was used. Contigs are indicated in magenta and blue. ESL0284 and ESL0368 are completely assembled genomes and were used as reference to sort the contigs of the other assemblies. The relatively few homologous regions between ESL0387 and ESL0378 reflects the deep divergence between the two different sub-lineages of Alpha2.2, i.e. the clade of *Bombella apis* (ESL0368, ESL0380, and ESL0387), and the clade of a novel species within the genus *Bombella* ( ESL0378 and ESL0385).

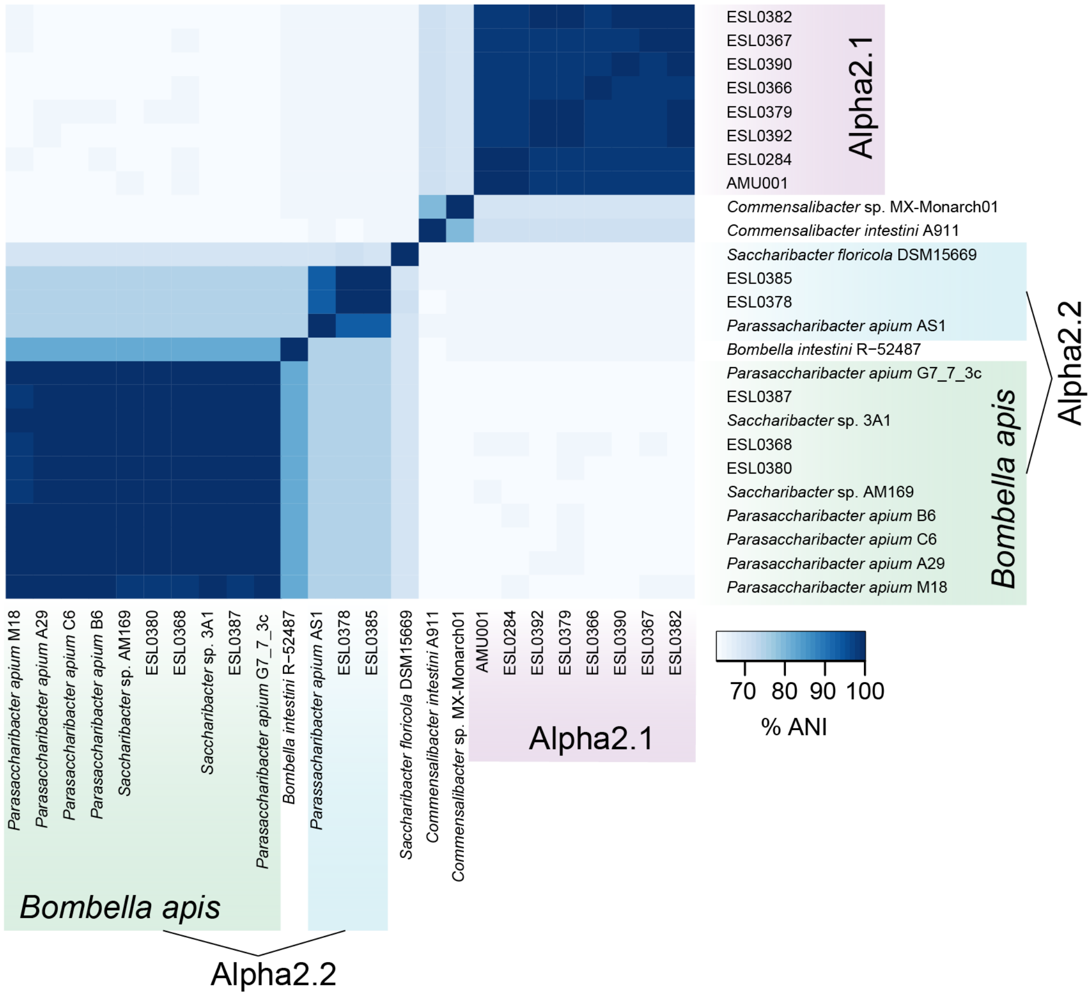


**Figure S2. Average nucleotide identity between strains of Alpha2.1, Alpha2.2, and closely related isolates**. Strains isolated from honey bees or their hive environment are highlighted in color. Alpha2.2 consists of two sub-lineages that share relatively low ANI, *Bombella apis* and a potentially new species within the genus *Bombella*.

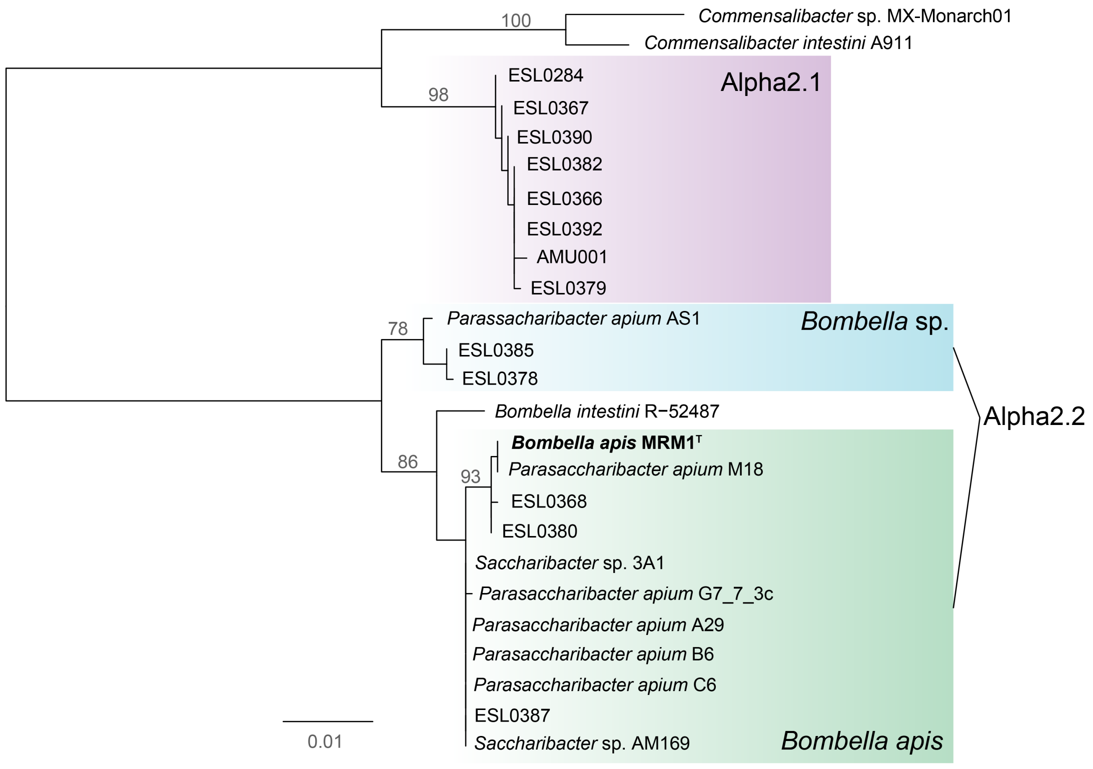


**Figure S3. Phylogeny of Alpha2.1 and Alpha2.2 based on 16S rRNA gene sequences**. The mafft (Katoh and Standley 2013) alignment compirses1,500 sites. The phylogeny was calculated using PHYML with the GTR model in Geneious. 100 bootstrap trees were calculated and nodes with a bootstrap support >80 are shown on branches.

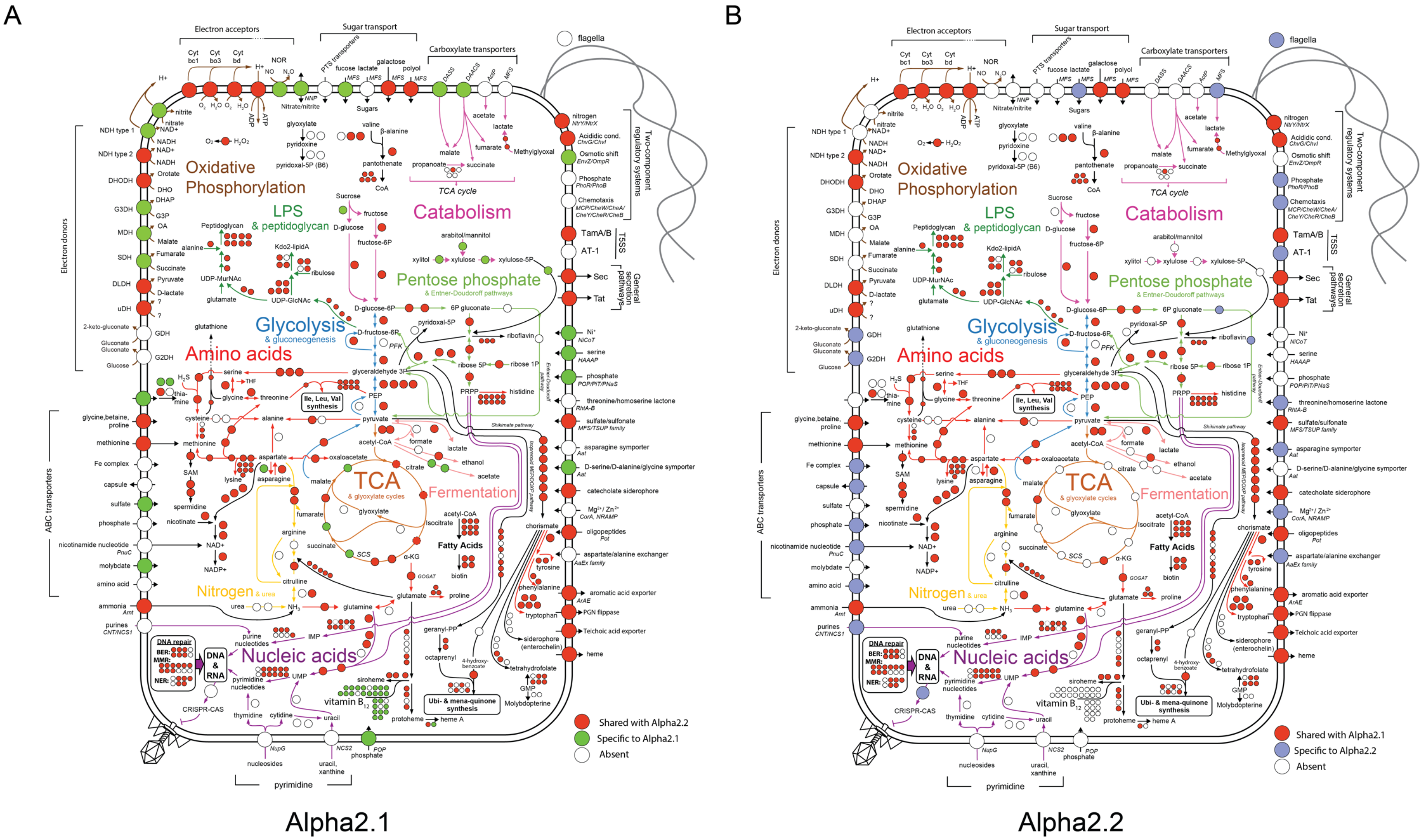


**Figure S4. Detailed metabolic maps of Alpha2.1 and Alpha2.2**. Red circles indicate shared core gene content. Green and purple indicate phylotype specific core gene content. Number of circles indicate number of enzymes needed for a given pathway. The blueprint of these metabolic maps was adapted from Kwong et al (2014).

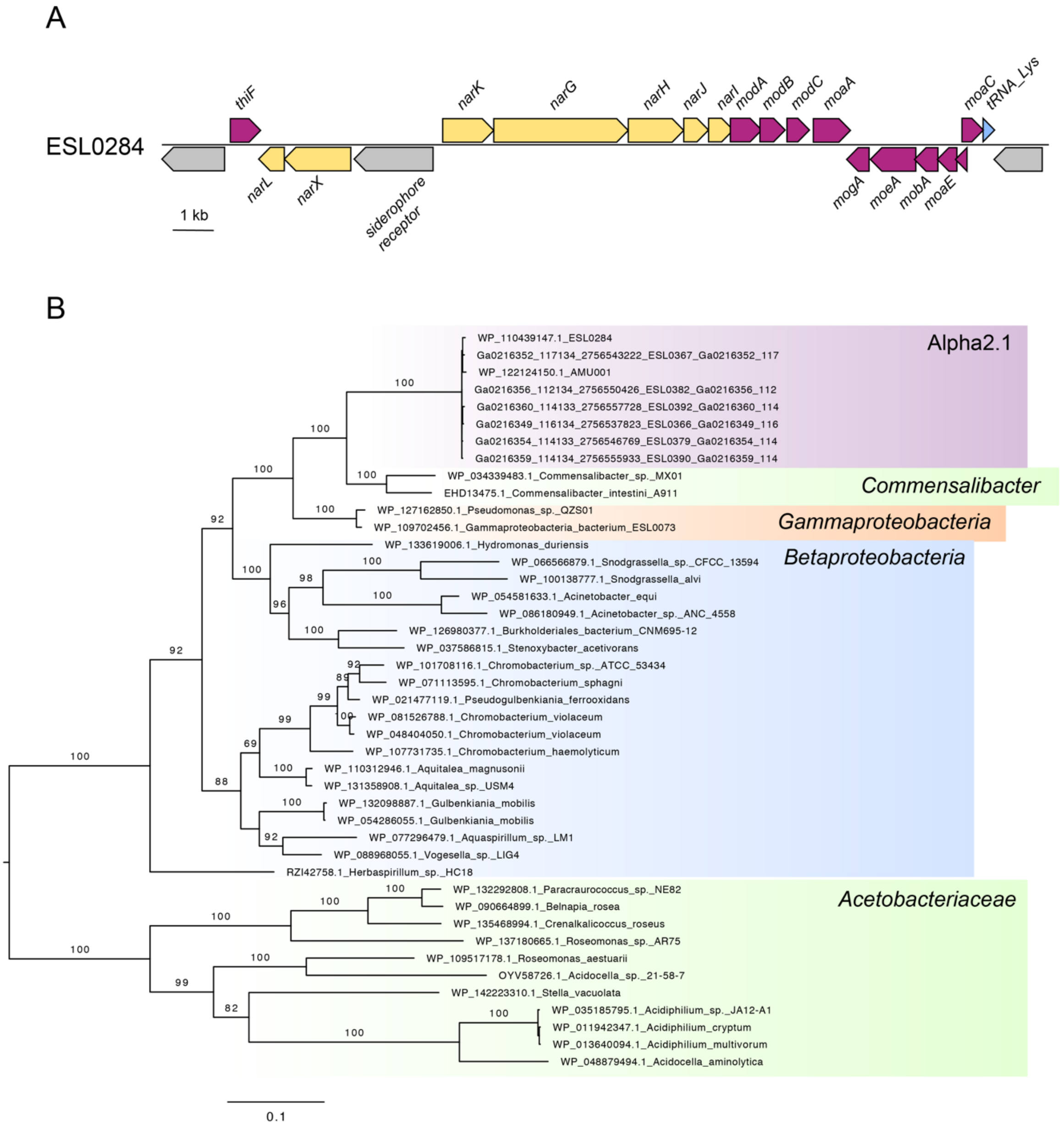


**Figure S5. Genomic regions encoding the nitrate reductase and molybdopterin biosynthesis genes of Alpha2.1 and gene tree of the alpha subunit of the nitrate reductase gene.** (A) Genomic locus of Alpha2.1 strain ESL0284 encoding the nitrate reductase genes (in yellow) and the biosynthesis genes for the co-factor molybdopterin (magenta). Other genes are shown in grey. A tRNA gene at the border of the genomic locus is shown in blue. Gene names are indicated if available. (B) Phylogenetic tree of the nitrate reductase alpha subunit gene (*narG*) of Alpha2.1 and related homologs found by BlastP against the nr database on NCBI. Distant homologs present in Acetobacteraceae were identified by searching with BlastP against the subset of 'Acetobacteraceae' in the nr database. A 1,269 sites-sites long alignment was produced with mafft mafft (Katoh and Standley 2013). The tree was inferred using RAXML with the PROTCATWAG model and 100 bootstrap replicates. Bootstrap values ≥80 are shown on branches.

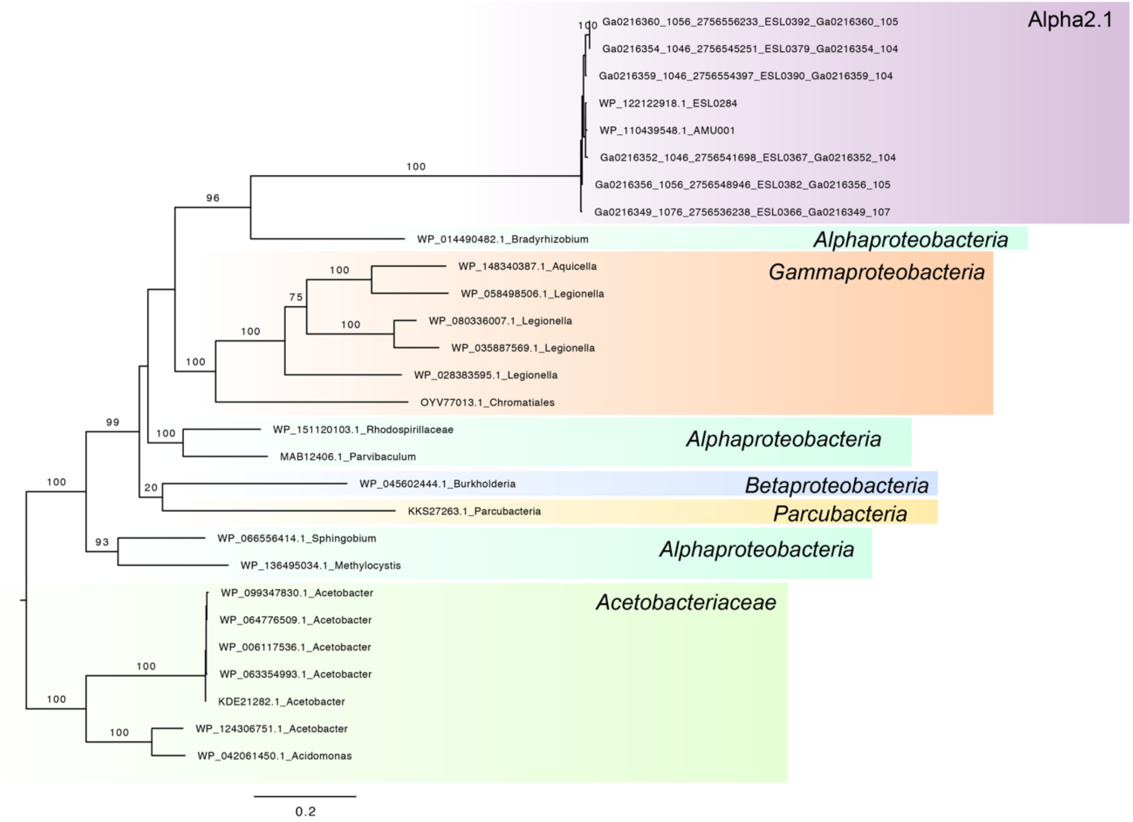


**Figure S6. Gene tree of the nitric oxide reductase of Alpha2.1.** The phylogenetic tree was inferred using RAXML with the PROTCATWAG model and 100 bootstrap replicates (Stamatakis 2014). Bootstrap values are depicted on the corresponding branch when ≥80. The most closely related homologs of the nitric oxide reductase of Alpha2.1 were found by BlastP against the nr database on NCBI. Distant homologs present in Acetobacteraceae were identified by searching with BlastP against the subset of ‘Acetobacteraceae’ in the nr database on NCBI. Sequences were aligned with mafft (Katoh and Standley 2013) resulting in 833 aligned amino acid sites.

#### Supplementary Tables

Table S1. Strain and genome feature table.

Table S2: ANI values.

Table S3: List of all gene families.

Table S4: List of the conserved pathways/functions between Alpha2.1 and Alpha2.2 listing corresponding gene families.

Table S5. List of shared and phylotype-specific core gene families as displayed in Figure 3A and B.

Table S6. Membrane-bound dehydrogenases involved in respiration.
